## Appendix 1 for "Spike frequency adaptation supports network computations on temporally dispersed information"

Darjan Salaj[1,†], Anand Subramoney[1,†], Ceca Kraišniković[1,†], Guillaume Bellec[1,2], Robert Legenstein[1], Wolfgang Maass[1,*]

June 18, 2021

[1] *Institute of Theoretical Computer Science, Graz University of Technology, Inffeldgasse 16b, Graz, Austria*
[2] *Laboratory of Computational Neuroscience, Ecole Polytechnique Fédérale de Lausanne (EPFL), Bâtiment AAB, offices 135-141, CH-1015 Lausanne*
[†] *These authors contributed equally to this work.*

#### 1 Autocorrelation based intrinsic time scale of neurons trained on STORE-RECALL task.

We wondered whether the adaptive firing threshold of LIF neurons with SFA affects the autocorrelation function of their firing activity — termed intrinsic time scale in (Wasmuht et al., 2018). We tested this for an SNN consisting of 200 LIF neurons without and 200 LIF neurons with SFA that was trained to solve a one-dimensional version of the STORE-RECALL task. It turned out that during the delay between STORE and RECALL these intrinsic time constants were in the same range as those measured in the monkey cortex, see Figure 1C in (Wasmuht et al., 2018). Furthermore, neurons of the trained SNN exhibited very similar distributions of these time constants (see Appendix figure 1), suggesting that these intrinsic time constants are determined largely by their network inputs, and less by the neuron type.

#### 2 sMNIST task with sparsely connected SNN obeying Dale's law

This task has originally been used as a temporal processing benchmark for ANNs, and has successfully been solved with the Long Short-Term Memory (LSTM) type of ANNs (Hochreiter and Schmidhuber, 1997). LSTM units store information in registers – like a digital computer – so that the stored information cannot be perturbed by ongoing network activity. Networks of LSTM units or variations of such units have been widely successful in temporal processing and reach the level of human performance for many temporal computing tasks.

Since LSTM networks also work well for tasks on larger time-scales, for comparing SNNs with LSTM networks, we used a version of the task with 2 ms presentation time per pixel, thereby doubling the length of sequences to be classified to 1568 ms. Grey values of pixels were presented to the LSTM network simply as analog values. A trial of a trained SNN with SFA (with an input sequence that encodes a handwritten digit "3" using population rate coding) is shown in Appendix figure 3B. The top row of Figure 3B shows a version where the grey value of the currently presented pixel is encoded by population coding, through the firing probability of 80 input neurons. Somewhat better performance was achieved when each of the 80 input neurons was associated with a particular threshold for the grey value, and this input neuron fired whenever the grey value crossed its threshold in the transition from the previous to the current pixel (this input convention was used to produce the results below).

Besides a fully connected network of LIF neurons with SFA, we also tested the performance of a variant of the model, called SC-SNN, that integrates additional constraints of SNNs in the brain: It is sparsely connected (12% of possible connections are present) and consists of 75% excitatory and 25% inhibitory neurons that adhere to Dale's law. By adapting the sparse connections with the rewiring method in (Bellec et al., 2018) during $BPTT$ training, the SC-SNN was able to perform even better than the fully-connected SNN of LIF neurons with SFA. The resulting architecture of the SC-SNN is shown in Appendix figure 3C. Its activity of excitatory and inhibitory neurons, as well as the time courses of adaptive thresholds for (excitatory) LIF neurons with SFA of the SC-SNN are shown in Appendix figure 3B. In this setup, the SFA had $\tau_a = 1400\,\mathrm{ms}$. When we used an SNN with SFA, we improved the accuracy on this task to 96.4% which approaches the accuracy of the artificial LSTM model which reached the accuracy of 98.0%.

We also trained a liquid state machine version of the SNN model with SFA where only the readout neurons are trained. This version of the network reached the accuracy of $63.24 \pm 1.48\%$ over 5 independent training runs.

### 3 Google Speech Commands

We trained SNNs with and without SFA on the keyword spotting task with Google Speech Commands Dataset (Warden, 2018) (v0.02). The dataset consists of 105,000 audio recordings of people saying thirty different words. Fully connected networks were trained to classify audio recordings, that were clipped to one second length, into one of 12 classes (10 keywords, as well as two special classes for silence and unknown words; the remaining 20 words had to be classified as "unknown"). Comparison of the maximum performance of trained spiking networks against state-of-the-art artificial recurrent networks is shown in Table 1. Averaging over 5 runs, the SNN with SFA reached $90.88 \pm 0.22\%$, and the SNN without SFA reached $88.79 \pm 0.16\%$ accuracy. Thus an SNN without SFA can already solve this task quite well, but the inclusion of SFA halves the performance gap to the published state-of-the-art in machine learning. The only other report on a solution to this task with spiking networks is (Zenke and Vogels, 2020). There the authors train a network of LIF neurons using surrogate gradients with $BPTT$ and achieve $85.3 \pm 0.3\%$ accuracy on the full 35 classes setup of the task. In this setup, the SNN with SFA reached $88.5 \pm 0.16\%$ test accuracy.

Features were extracted from the raw audio using the Mel Frequency Cepstral Coefficient (MFCC) method with 30 ms window size, 1 ms stride, and 40 output features. The network models were trained to classify the input features into one of the 10 keywords (yes, no, up, down, left, right, on, off, stop, go) or to two special classes for silence or unknown word (where the remainder of 20 recorded keywords are grouped). The training, validation and test set were assigned 80, 10, and 10 percent of data respectively while making sure that audio clips from the same person stay in the same set.

All networks were trained for 18,000 iterations using the Adam optimizer with batch size 100. The output spikes of the networks were averaged over time, and the linear readout layer was applied to those values. During the first 15,000 iterations, we used a learning rate of 0.001 and for the last 3000, we used a learning rate of 0.0001. The loss function contained a regularization term (scaled with coefficient 0.001) that minimizes the squared difference of average firing rate between individual neurons and a target firing rate of 10 Hz.

Both SNNs with and without SFA consisted of 2048 fully connected neurons in a single recurrent layer. The neurons had a membrane time constant of $\tau_m = 20\,\mathrm{ms}$, the adaptation time constant of SFA was $\tau_a = 100\,\mathrm{ms}$, adaptation strength was $\beta = 2\,\mathrm{mV}$. The baseline threshold was $v_{\mathrm{th}} = 10\,\mathrm{mV}$, and the refractory period was $2\,\mathrm{ms}$. The synaptic delay was $1\,\mathrm{ms}$.

### 4 Delayed-memory XOR

We also tested the performance of SNNs with SFA on a previously considered benchmark task, where two items in the working memory have to be combined non-linearly: the Delayed-memory

XOR task (Huh and Sejnowski, 2018). The network is required to compute the exclusive-or operation on the history of input pulses when prompted by a go-cue signal, see Appendix figure 4.

The network received on one input channel two types of pulses (up or down), and a go-cue on another channel. If the network received two input pulses since the last go-cue signal, it should generate the output "1" during the next go-cue if the input pulses were different or "0" if the input pulses were the same. Otherwise, if the network only received one input pulse since the last go-cue signal, it should generate a null output (no output pulse). Variable time delays are introduced between the input and go-cue pulses. The time scale of the task was 600 ms which limited the delay between input pulses to 200 ms.

This task was solved in (Huh and Sejnowski, 2018), without providing performance statistics, by using a type of neuron that has not been documented in biology — a non-leaky quadratic integrate and fire neuron. We are not aware of previous solutions by networks of LIF neurons. To compare and investigate the impact of SFA on network performance in the delayed-memory XOR task, we trained SNNs, with and without SFA, of the same size as in (Huh and Sejnowski, 2018) — 80 neurons. Across 10 runs, SNNs with SFA solved the task with $95.19 \pm 0.014\%$ accuracy, whereas the SNNs without SFA converged at lower $61.30 \pm 0.029\%$ accuracy.

The pulses on the two input channels were generated with 30 ms duration and the shape of a normal probability density function normalized in the range $[0, 1]$. The pulses were added or subtracted from the baseline zero input current at appropriate delays. The go-cue was always a positive current pulse. The 6 possible configurations of the input pulses ($+$, $-$, $++$, $--$, $+-$, $-+$) were sampled with equal probability during training and testing.

Networks were trained for 2000 iterations using the Adam optimizer with batch size 256. The initial learning rate was 0.01 and every 200 iterations the learning rate was decayed by a factor of 0.8. The loss function contained a regularization term (scaled with coefficient 50) that minimizes the squared difference of the average firing rate of individual neurons from a target firing rate of 10 Hz. This regularization resulted in networks with a mean firing rate of 10 Hz where firing rates of individual neurons were spread in the range [1, 16] Hz.

Both SNNs with and without SFA consisted of 80 fully connected neurons in a single recurrent layer. The neurons had a membrane time constant of $\tau_m = 20\,\mathrm{ms}$, a baseline threshold $v_{\mathrm{th}} = 10\,\mathrm{mV}$, and a refractory period of 3 ms. SFA had an adaptation time constant of $\tau_a = 500\,\mathrm{ms}$ and an adaptation strength of $\beta = 1\,\mathrm{mV}$. The synaptic delay was 1 ms. For training the network to classify the input into one of the three classes, we used the cross-entropy loss between the labels and the softmax of three linear readout neurons. The input to the linear readout neurons were the neuron traces that were calculated by passing all the network spikes through a low-pass filter with a time constant of 20 ms.

### 5 12AX task in a noisy network

As a control experiment, aimed at testing the robustness of the solution (performance as a function of the strength of added noise), we simulated the injection of an additional noise current into all LIF neurons (with and without SFA). The previously trained network (trained without noise) was reused and tested on a test set of $2,000$ episodes. In each discrete time step, the noise was added to the input current $I_j(t)$ (see equation (4) in the main text), hence affecting the voltage of the neuron:

$$I_j(t) \quad = \quad \sum_i W_{ji}^{\mathrm{in}} x_i(t - d_{ji}^{\mathrm{in}}) + \sum_i W_{ji}^{\mathrm{rec}} z_i(t - d_{ji}^{\mathrm{rec}}) + I_{\mathrm{noise}}, \tag{1}$$

where $I_{\mathrm{noise}}$ was drawn from a normal distribution with mean zero, and standard deviation $\sigma \in \{0.05, 0.075, 0.1, 0.2, 0.5\}$.

Performance of the network without noise was 97.85% (performance of one initialization of the network with 100 LIF neurons with SFA and 100 LIF neurons without SFA). During testing, including the noise current of mean zero and standard deviation $\sigma \in \{0.05, 0.075, 0.1, 0.2, 0.5\}$ lead to the performance of 92.65%, 89.05%, 80.25%, 27.25%, 0.25%, respectively. The network performance degrades gracefully up to a current of standard deviation of about 0.1.

For an illustration of the effect of noise, see Appendix figure 5 and 6. There, we compare the output spikes, adaptive threshold, and membrane voltage of one neuron with noise current to the versions without noise. The shown simulations started from exactly the same initial condition and noise with standard deviation 0.05 (0.075) was injected only into the shown neuron (other neurons did not receive any noise current). One sees that even this weak noise current produces a substantial perturbation of the voltage, adaptive threshold, and spiking output of the neuron.

### 6 Duplication/Reversal task

A zoom-in for the rasters shown in Figure 4A (from the main text) is shown in Appendix figure 7, for the time period $3-4$ seconds.

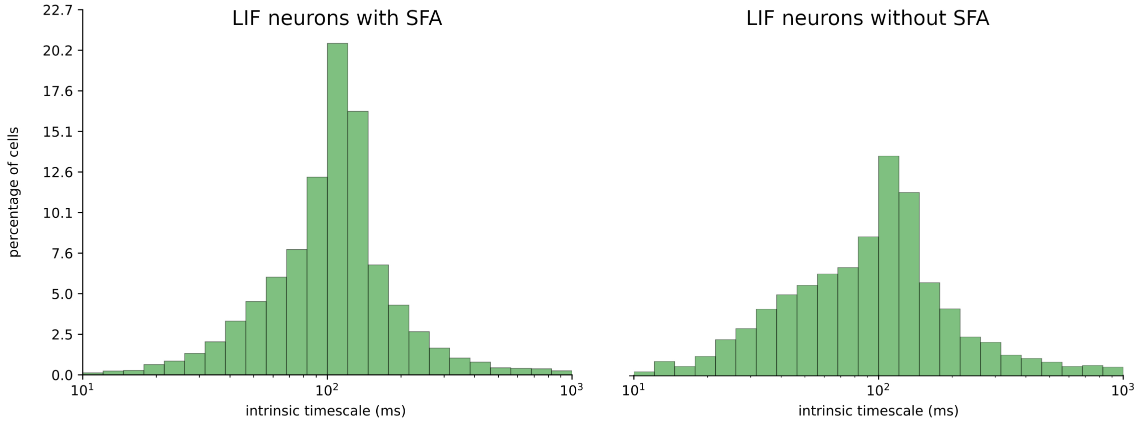


**Appendix figure 1: Histogram of the intrinsic time scale of neurons trained on STORE-RECALL task.** We trained 64 randomly initialized SNNs consisting of 200 LIF neurons with and 200 without SFA on the single-feature STORE-RECALL task. Measurements of the intrinsic time scale were performed according to (Wasmuht et al., 2018) on the spiking data of SNNs solving the task after training. Averaged data of all 64 runs is presented in the histogram. The distribution is very similar for neurons with and without SFA.

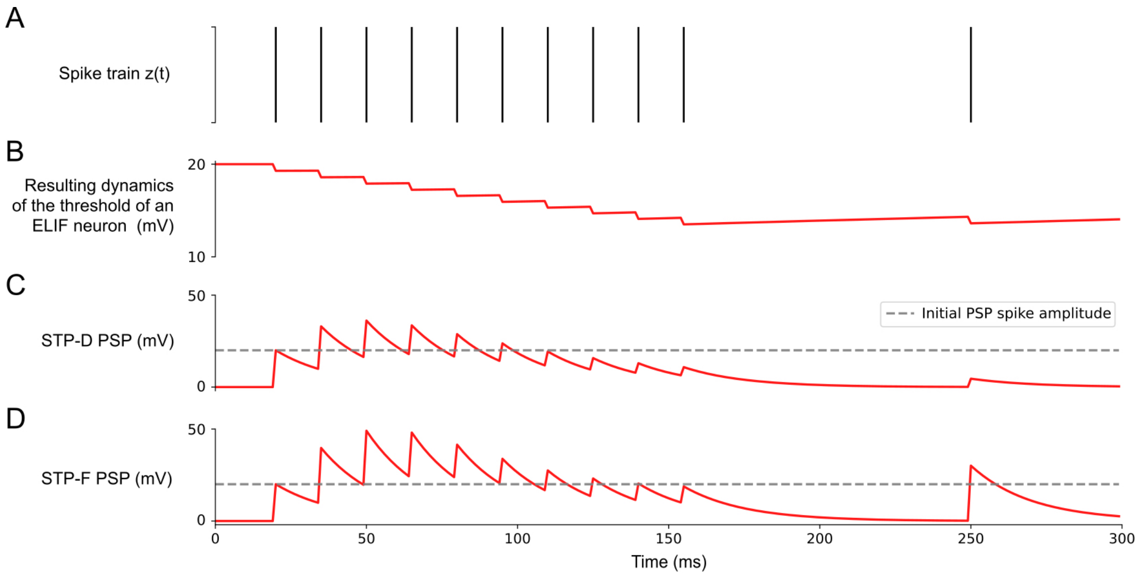


**Appendix figure 2: Illustration of models for an inversely adapting ELIF neuron, and for short-term synaptic plasticity.** (**A**) Sample spike train. (**B**) The resulting evolution of firing threshold for an inversely adapting neuron (ELIF neuron). (**C-D**) The resulting evolution of the amplitude of postsynaptic potentials (PSPs) for spikes of the presynaptic neuron for the case of a depression-dominant (STP-D: D$\gg$F) and a facilitation-dominant (STP-F: F$\gg$D) short-term synaptic plasticity.

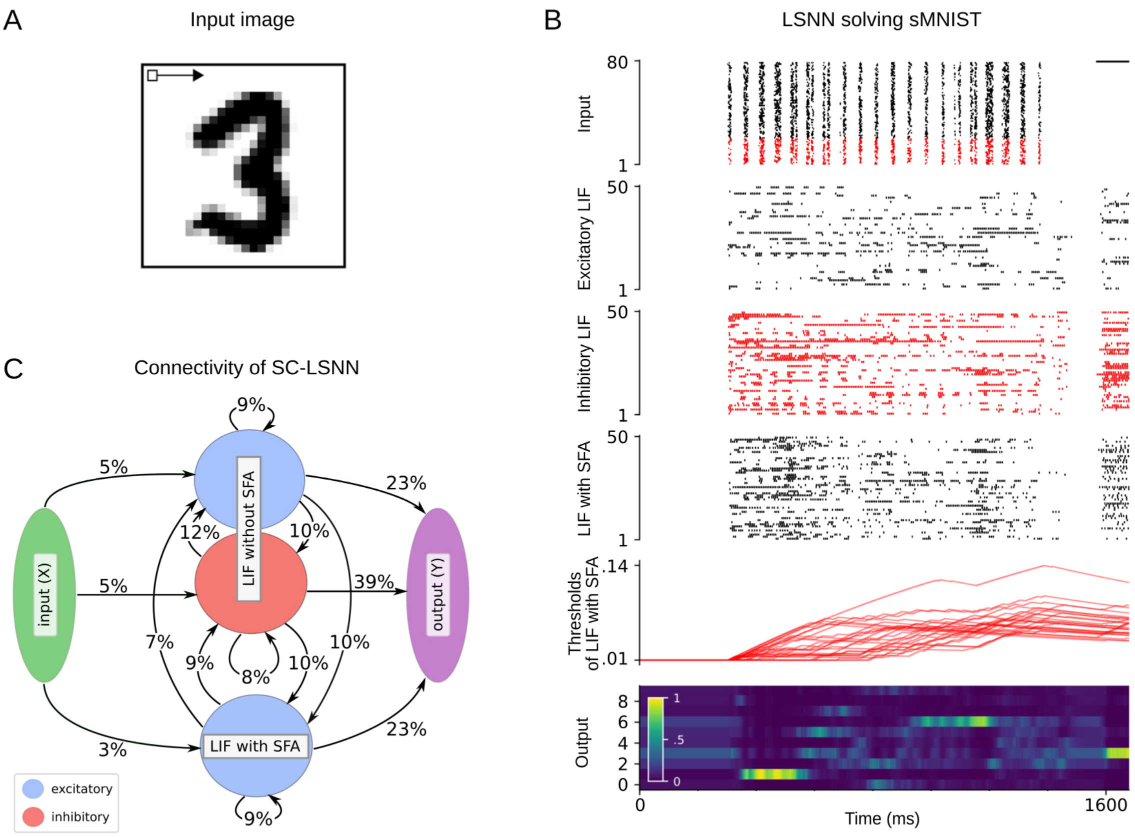


**Appendix figure 3: sMNIST time series classification benchmark task.** (**A**) Illustration of the pixel-wise input presentation of handwritten digits for sMNIST. (**B**) Rows top to bottom: Input encoding for an instance of the sMNIST task, network activity, and temporal evolution of firing thresholds for randomly chosen subsets of neurons in the SC-SNN, where 25% of the LIF neurons were inhibitory (their spikes are marked in red). The light color of the readout neuron for digit "3" around 1600 ms indicates that this input was correctly classified. (**C**) Resulting connectivity graph between neuron populations of an SC-SNN after BPTT optimization with DEEP R on sMNIST task with 12% global connectivity limit.

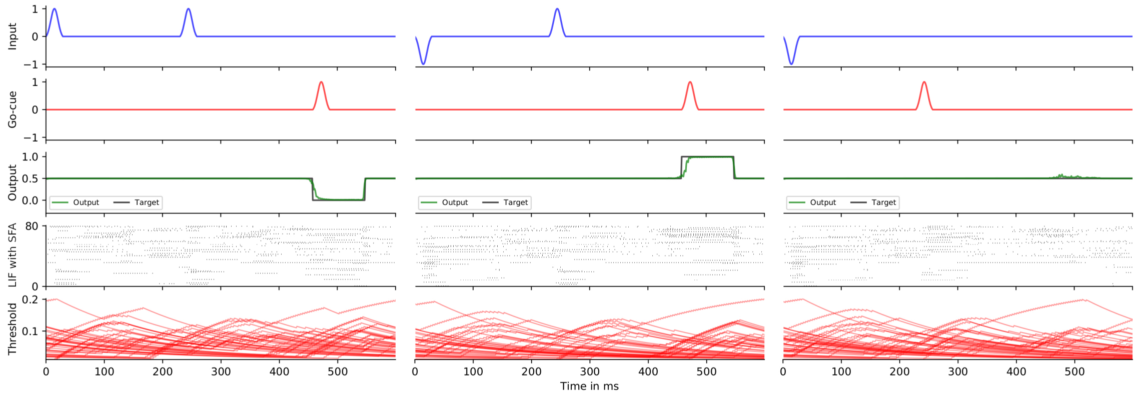


**Appendix figure 4: Delayed-memory XOR task.** Rows top to bottom: Input signal, Go-cue signal, network readout, network activity, and temporal evolution of firing thresholds.

| Model | test accuracy (%) |
|---|---|
| FastGRNN-LSQ (Kusupati et al., 2018) | 93.18 |
| SNN with SFA | 91.21 |
| SNN | 89.04 |

**Appendix table 1: Google Speech Commands.** Accuracy of the spiking network models on the test set compared to the state-of-the-art artificial recurrent model reported in (Kusupati et al., 2018). Accuracy of the best out of 5 simulations for SNNs is reported.

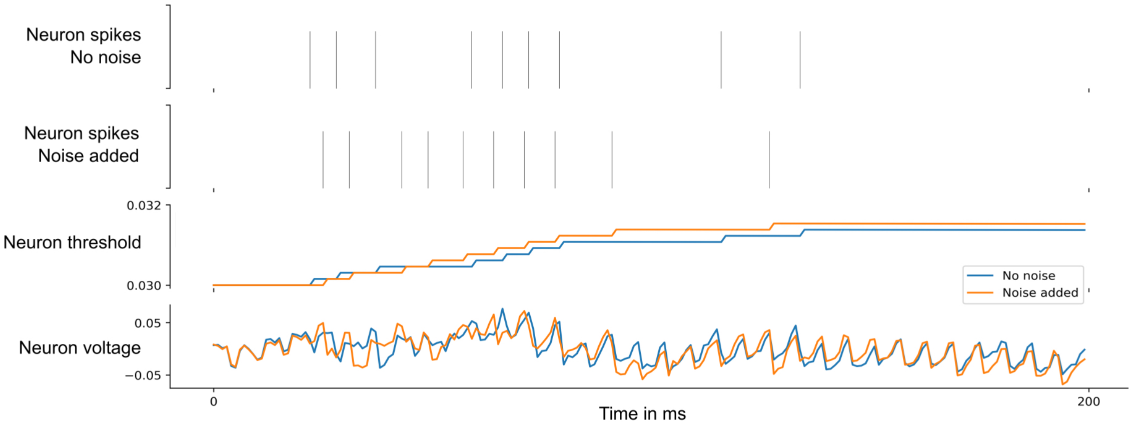


**Appendix figure 5: Effect of a noise current with zero mean and standard deviation 0.05 added to a single neuron in the 12AX task.** Spike train of a single neuron without noise, followed by spike train in the presence of the noise, adaptive threshold of the neuron that corresponds to the spike train with no noise (shown in blue), spike train with noise present (shown in orange), and corresponding neuron voltages over the time course of 200 ms.

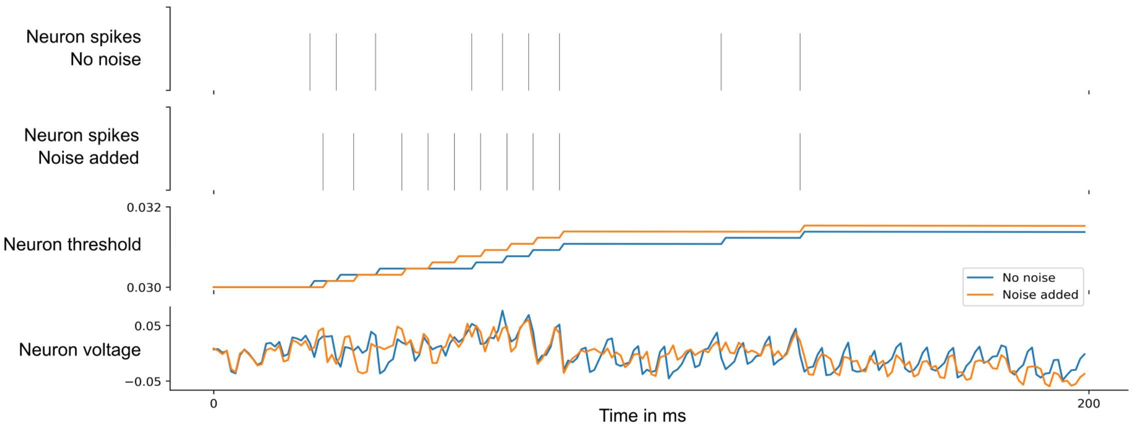


**Appendix figure 6: Effect of a noise current with zero mean and standard deviation 0.075 added to a single neuron in the network for the 12AX task.** Spike train of a single neuron without noise, followed by spike train in the presence of the noise, adaptive threshold of the neuron that corresponds to the spike train with no noise (shown in blue), spike train with noise present (shown in orange), and corresponding neuron voltages over the time course of 200 ms.

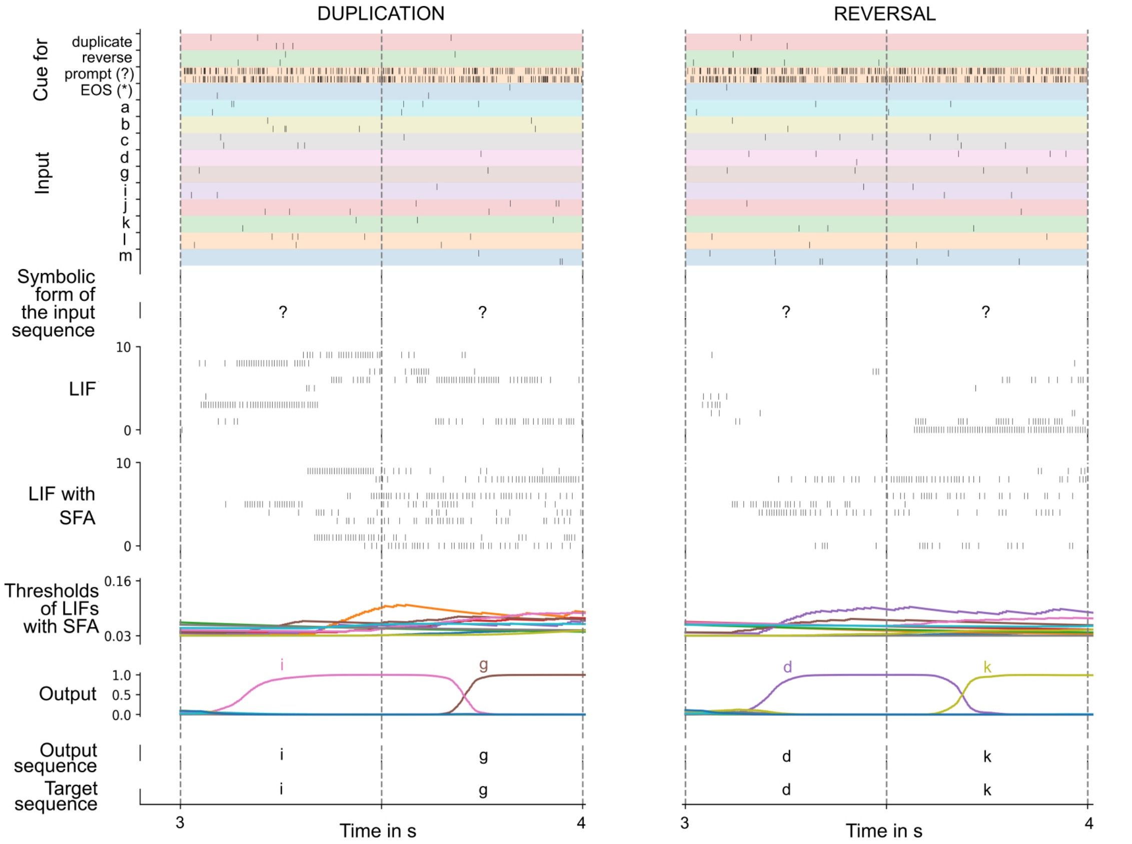


**Appendix figure 7: A zoom-in of the spike raster for a trial solving Duplication task (left) and Reversal task (right).** A sample episode where the network carried out sequence duplication (left) and sequence reversal (right), shown for the time period of 3 - 4 ms (2 steps after the start of network output). Top to bottom: Spike inputs to the network (subset), sequence of symbols they encode, spike activity of 10 sample LIF neurons (without and with SFA) in the SNN, firing threshold dynamics for these 10 LIF neurons with SFA, activation of linear readout neurons, output sequence produced by applying argmax to them, target output sequence.

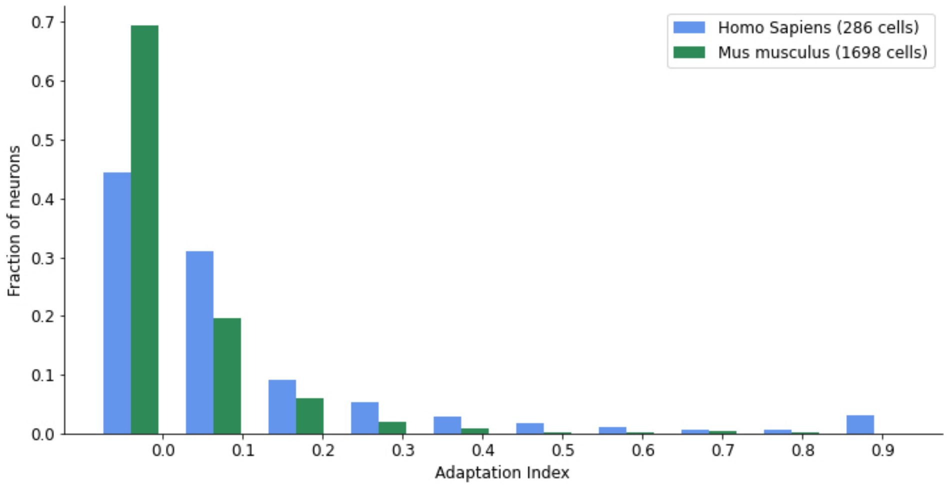


**Appendix figure 8: Distribution of adaptation index from Allen Institute cell measurements (Allen Institute, 2018).**
